# The influence of stereopsis on visual saliency in a proto-object based model of selective attention

**DOI:** 10.1101/2022.04.04.486976

**Authors:** Takeshi Uejima, Elena Mancinelli, Ernst Niebur, Ralph Etienne-Cummings

## Abstract

Some animals including humans use stereoscopic vision which reconstructs spatial information about the environment from the disparity between images captured by eyes in two separate adjacent locations. Like other sensory information, such stereoscopic information is expected to influence attentional selection. We develop a biologically plausible model of binocular vision to study its effect on bottom-up visual attention, i.e., visual saliency. In our model, the scene is organized in terms of proto-objects on which attention acts, rather than on unbound sets of elementary features. We show that taking into account the stereoscopic information improves the performance of the model in the prediction of human eye movements with statistically significant differences.

## 1 Introduction

We are surrounded by three-dimensional space; however, each retina captures only a two-dimensional image. Each retinal image individually can contain clues for depth information such as shading, looming sizes, and occlusion, with the latter including the presence of T-junctions (Nakayama et al., 1995; von der Heydt, 2015; Welchman, 2016). In addition, binocular vision which uses triangulation by two eyeballs, provides a reliable cue for depth and results in vivid three-dimensional (3D) perception. Not only is this capability used for range finding, some animals with front-facing eyes, including humans, also exploit binocular stereopsis for camouflage breaking (Nityananda & Read, 2017); in the words of Bela Julesz, “with stereoscopic vision there is no camouflage” (Julesz, 1989). This is likely due to the organization of visual scenes into (proto-)objects: camouflage exploits erroneous assignment of perceptual edges which is made more difficult by the existence of explicit depth differences at object borders, and their absence within object borders (Adams et al., 2019; Poggio & Poggio, 1984)

Binocular vision has been studied both in neuroscience and in engineering, the former primarily focusing on revealing how nervous systems achieve stereovision and the latter on finding efficient and precise algorithms. In both fields, a major difficulty is the stereo correspondence problem: to find out which features in two retinal images originate from the same point in 3D space. The problem may seem trivial because we effortlessly and quickly solve it in daily life. Nonetheless, it is not simple, and the brain devotes multiple cortical areas to solve it (Cumming & DeAngelis, 2001; Kumano et al., 2008; Tanabe et al., 2004). The mechanisms processed in the primary visual cortex have been extensively studied, and the “disparity energy model” is widely accepted as it agrees well with data from neurophysiological experiments (Ohzawa et al., 1990, 1997).

Binocular information, together with other visual features, not only underlies functions like object recognition but presumably also provides inputs for the determination of which parts of the visual scene are the most relevant, i.e. which require detailed processing. Identifying these regions is the task solved by visual selective attention. In general, this is a highly complex function which involves perceptual and cognitive processes at many levels. An important part of this function is data-driven, or bottom-up attention, that finds the most relevant image regions based on low-level visual features and their combinations. These regions are usually called the most “salient” areas of the scene. The seminal work by Koch and Ullman (Koch & Ullman, 1985) elaborated a systematic way to find these regions in the form of a saliency map which ranks the level of saliency at different locations in the visual scene. Predictions of this theory need to be compared with behavioral observations. Two considerations are of relevance here. First, the saliency map was originally developed for covert attention whose effects may not result in overt attention, i.e. eye movements. There are methods to measure the effects of covert visual attention, e.g. (Posner, 1980) but in practice, it is much easier to measure the state of overt attention (Parkhurst et al., 2002) which is known to correlate with covert attention (Deubel & Schneider, 1996; Hoffman & Subramaniam, 1995; Moore & Fallah, 2001; Van der Stigchel & Theeuwes, 2007). Second, the saliency map only takes account of bottom-up information. To minimize (though not eliminate) the effects of top-down attention, which is goal-directed and depends on the internal state of the observer in addition to the visual input, (Parkhurst et al., 2002) and later studies use a free-viewing paradigm.

Many models of visual saliency rely on local contrast in low-level features such as intensity, color, and orientation. However, a plethora of studies in psychology and neurophysiology have shown that visual attention is also influenced by the rapid perceptual organization of the visual scene into tentative objects rather than the basic features themselves (Egly et al., 1994; Einhäuser et al., 2008; Nuthmann & Henderson, 2010; Qiu et al., 2007; Stoll et al., 2015; Zhou et al., 2000). These tentative objects or areas that possess “objectness” are called proto-objects (Rensink, 2000). A proto-object based saliency model was shown to predict eye fixations with good accuracy (Russell et al., 2014). While originally this model used information from maps of intensity, color, and orientation, it was later extended to additionally utilize motion, depth, and texture features (Hu et al., 2016; Mancinelli et al., 2018; Molin et al., 2015; Uejima et al., 2020), and also implemented in biofidelic neuromorphic hardware (Iacono et al., 2019).

In this paper, we propose a model of biological stereopsis and incorporate it into that proto-object based saliency model. While depth and disparity features have been integrated into the model previously (Hu et al., 2016; Mancinelli et al., 2018), we discuss below why our approach employs a different mechanism to exploit binocular disparity that is biologically plausible. As in previous work, the output of our model is a saliency map, and we compare it with published human fixation data obtained while participants freely viewed stereogram images. As we show below, our model shows better predictive performance than the original two-dimensional (2D) proto-object based saliency model.

## 2 Related Studies

### 2.1 Stereopsis and Eye Fixations

Many studies have sought an understanding of how the brain achieves stereoscopic vision. Since Julesz introduced random dot stereograms (RDSs) (Julesz, 1971), which do not include any 2D depth cues and provide only disparity information, neuroscientists have used these stimuli to study brain activity corresponding to depth perception solely generated by binocular disparity (Poggio et al., 1985). These experiments have also shown that no prior knowledge about objects is needed for stereo correspondence because an observer cannot perceive any nontrivial imaged contents before fusing the stereogram.

One of the major difficulties of stereoscopic vision is the correspondence problem, i.e. to find corresponding features in the two 2D images. A cooperative process is an early model to solve this problem, by finding matching points using iterative computations which minimize an error-measure (Marr, 1982; Marr & Poggio, 1979). While this could be realized in the nervous system, the biological system seems to process stereovision more rapidly than is expected from an iterative process, at least in its early stages. Neurophysiological studies have revealed that stereoscopic vision can be explained by the so-called disparity energy model, in which binocular simple cells sum the activity of monocular simple cells linearly. Subsequently, binocular complex cells sum the squared responses of quadrature pairs of the simple cells (Ohzawa, 1998; Ohzawa et al., 1990, 1997). Marr and Poggio pointed out that combining multi-spatial-frequency filters (i.e., receptive fields of cells) aids to prevent false matches between parts of one image to non-corresponding portions of the other image (Marr & Poggio, 1979). Their original idea employs a sequential coarse-to-fine structure that first computes coarse (low spatial frequencies) disparities and then proceeds to finer scales. Later, pooling multi-spatial-frequency features to find correspondence points based on information from all scales at once (rather than sequentially at different scales) was proposed (Fleet et al., 1996). While it is not clear whether the biological system employs a sequential mechanism or a simultaneous pooling algorithm, integration over multiple spatial-frequencies has been observed in primate area V4 (Kumano et al., 2008) and in primary visual cortex of cat (Baba et al., 2015), and we adopt it in our model.

Visual saliency has been widely studied in 2D (Bruce & Tsotsos, 2005; Hou et al., 2012; Itti et al., 1998; Itti & Koch, 2000; Judd et al., 2009; Koch & Ullman, 1985; Niebur & Koch, 1996), see (Borji et al., 2013) for a comparative study. While these models approach the problem from a mechanistic point of view, deep learning based models have also been used, and showed remarkable performance in predicting human fixations (Cornia et al., 2016; Huang et al., 2015; Kruthiventi et al., 2015; Kümmerer et al., 2014, 2016; Vig et al., 2014). Recently, some studies attempted to incorporate psychological concepts, Gestalt principles, into saliency models (Russell et al., 2014; Zhang & Sclaroff, 2016, 2013). Gestalt psychology argues that the whole of an object is more important than individual features for perception. In late years, this assertion has been supported by neurophysiological studies as figure-ground organization coding in visual cortex (Craft et al., 2007; Qiu & von der Heydt, 2005), which link perception and neural responses.

Although substantial efforts have been expended to model overt attention, the effect and mechanism of stereovision on saliency have not been fully examined. Reports investigating how depth information affects human eye movements (Gautier & Le Meur, 2012; Huynh-Thu & Schiatti, 2011; Jansen et al., 2009; Khaustova et al., 2013; Lang et al., 2012) showed that humans tend to fixate similar locations in situations with or without binocular information. More specifically, the fixation locations are almost the same for 3D and 2D images in long observation windows (20 seconds) but different in short observation windows (about four or five seconds) (Gautier & Le Meur, 2012; Jansen et al., 2009; Khaustova et al., 2013). Notably, researchers reported a tendency for humans to look at closer points soon after they look at an image (Gautier & Le Meur, 2012; Jansen et al., 2009; Lang et al., 2012).

Based on these behavioral results, researchers have proposed visual saliency models for 3D still scenes. Lang et al. calculated “depth priors” that indicates how eye fixations of human observers differed between 2D images and the corresponding 3D scenes, and they proposed to include these priors in existing 2D saliency models (Lang et al., 2012). Wang et al. took a Bayesian approach to incorporate depth effects on fixations (Wang et al., 2013), in which the parameters for the probability distribution were tuned by observed fixation data. Ma and Hang (Ma & Hang, 2015) proposed another learning-based model that employed a similar method for the Judd et al. model (Judd et al., 2009)for 2D static images, which includes various features including a face detection mechanism.

They extended the model to incorporate features from a depth map. In these data-driven approaches, the parameters are determined by human behavioral data and did not explicitly implement biological mechanisms. In this study, we instead implement algorithms inspired by the information processing principles employed in the primate brain. More specifically, we use the framework of a proto-object based model of perceptual organization and attentional control (Russell et al., 2014) and integrate a biologically-plausible stereovision mechanism in this model.

To our knowledge, to date there exist only two proto-object based saliency models that include depth features. The Hu et al. model takes a depth map as input along with a 2D image, which means depth information must be calculated or measured before it is used in the model (Hu et al., 2016). The depth map is then treated in a similar way to any other feature maps, e.g., the intensity map. The other model, proposed by Mancinelli et al. (Mancinelli et al., 2018) takes two images from the right and left cameras rather than a depth map. This approach is more biofidelic because the visual cortex is not provided with an explicit depth map as input, but instead, with the output from two retinae. However, the images in the Mancinelli et al model need to be rectified which either requires precise knowledge of the optical geometry, which often is not available, or additional knowledge of the depth at several locations in the scene. This is the main disadvantage of the model since it leaves open where this information comes from. We therefore take a different approach for modelling visual saliency based on Gestalt principles with a stereopsis mechanism that does not suffer from these limitations, by only requiring input from two cameras.

### 2.2 3D Eye Fixations Datasets

As mentioned previously, a widely accepted method to evaluate the quality of saliency models is to compare how well they can predict human eye fixations. Although many datasets of human fixations for 2D scenes have been published, only few are available for 3D stimuli.

The Gaze-3D dataset is a publicly-available 3D fixation dataset (Wang et al., 2013). It consists of 18 stereoscopic images and corresponding disparity and depth maps calculated by an optical flow method. The fixation data were collected from 35 participants sitting at 93 cm distance from a 26-inch display with a resolution of 1920×1200 pixels. Their eye tracking data were recorded from the left eye, meaning the fixation locations correspond to the left image.

The NCTU-3D dataset consists of 475 stereoscopic scenes and corresponding depth maps (Ma & Hang, 2015). The eye-tracking data were collected from 16 subjects. The 3D images were displayed on a 23-inch monitor with 1920×1080 pixels resolution, placed at 78.5 cm from the observers. Fig. **3** (a) shows an example of stimuli, fixation map, and depth map provided by the NCTU-3D dataset. The provided fixation data are based on right-eye tracking.

**Fig. 1.**
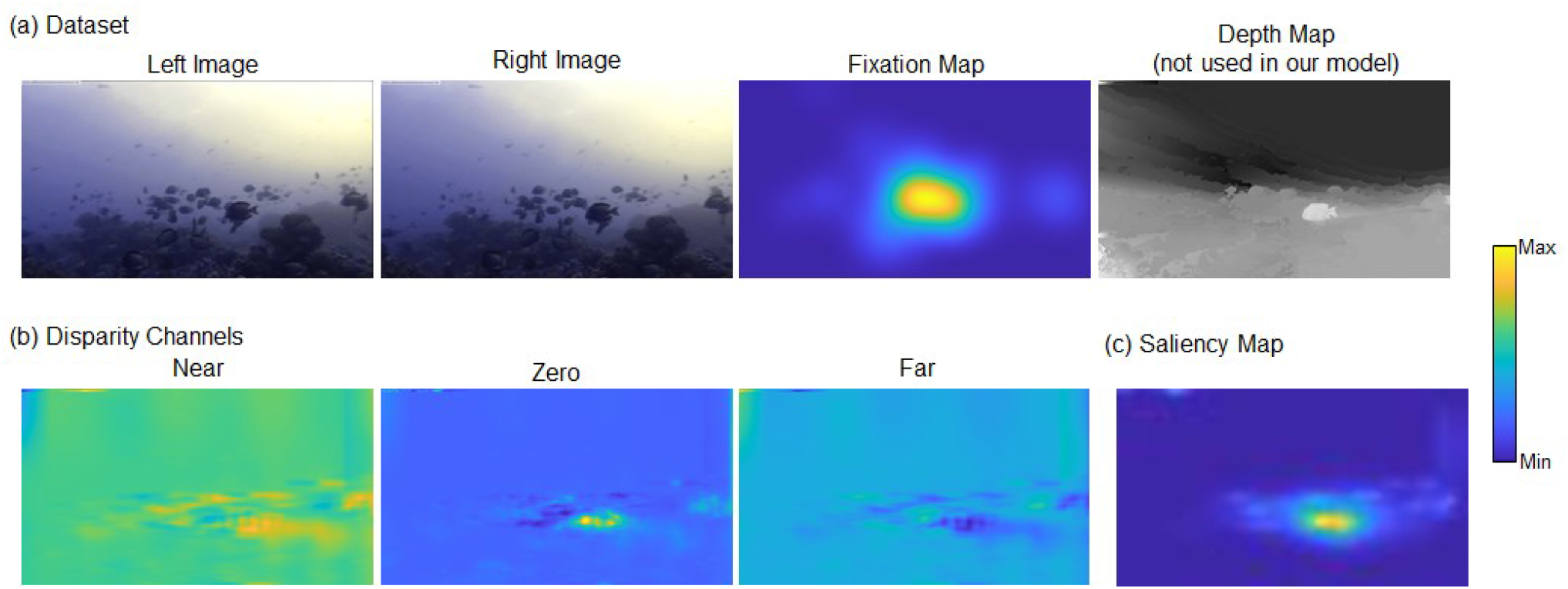
Examples of a 3D eye tracking dataset and the saliency map generated by the proposed model. (a) Stereo image, fixation map and depth map from the NCTU-3D dataset. The depth map is not used in our model. (b) Disparity channels of Near, Zero, and Far calculated by our model. A fish in the lower-right is present in the Near and Zero channels. (c) Saliency map generated from the disparity channels.

**Fig. 2.**
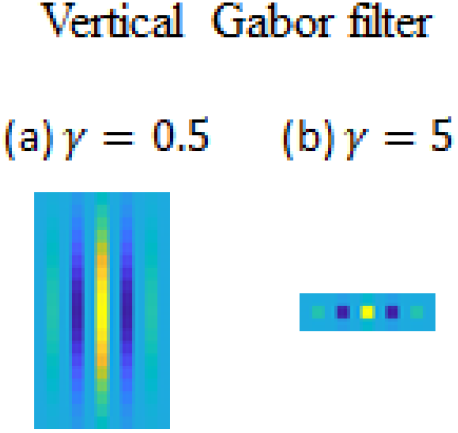
Vertical Gabor filters. (a) A high-aspect ratio Gabor filter. (b) A low-aspect ratio Gabor filter.

**Fig. 3.**
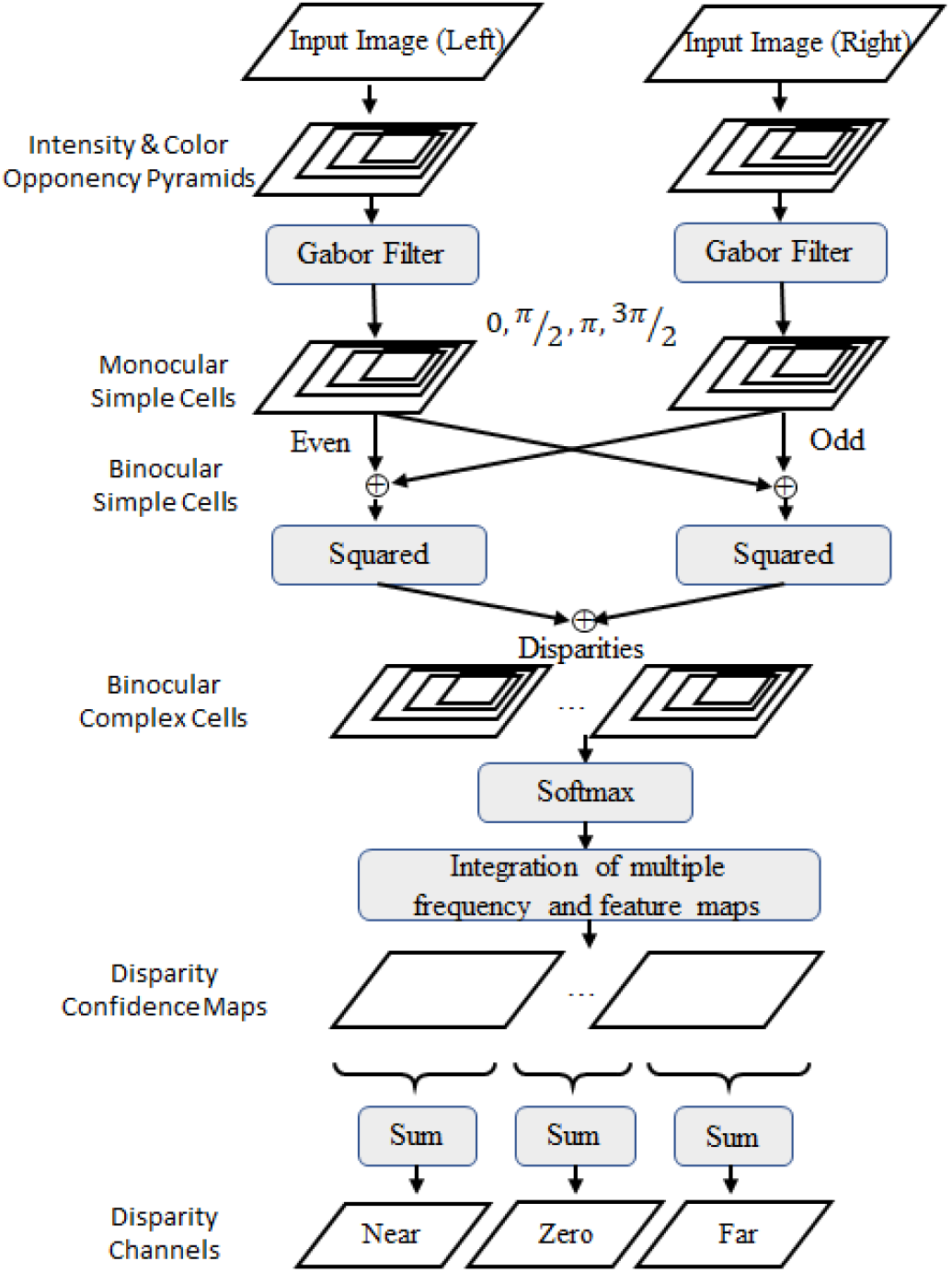
Schematic of disparity channel computation.

## 3 Proto-object based saliency model

### 3.1 Model framework

The disparity channels to be described in Section 3.2 provide input to a variation of the proto-object based saliency model which was originally introduced by Russell et al. (Russell et al., 2014). We use an improved algorithm developed by Uejima et al. (Uejima et al., 2020) but omit the texture features introduced in that model. Since the model framework used in this paper is the same as that in those prior studies, we explain it only briefly.

The model starts by extracting intensity, color, and orientation feature maps from two 2D images. Each input channel is processed by surface and edge detectors. The surface features are captured by center-surround receptive fields which are modeled as two 2D Gaussian filters. The edge detectors consist of Gabor filters modelled as simple cells, akin to those in 3.2.

The surface and edge maps are used to calculate border-ownership coding which is mainly observed in cortical area V2 (Zhou et al., 2000). A proto-object map is computed from the border-ownership maps by summing over orientation selective filters arranged tangentially to an annular region (Craft et al., 2007; Russell et al., 2014), and a conspicuity map is generated by normalizing the proto-object maps. The sum over proto-object maps of all features constitutes the final saliency map. In this paper, we call the model without the disparity channels (i.e., calculated from only intensity, color, and orientation features) the 2D model, and models that include the disparity channels 3D models (we define four such models, see below). The saliency maps of the 2D model can be computed from either the right or the left images. We use the image from the same side as that in the provided fixation map: left images for the Gaze-3D dataset and right images for the NCTU-3D dataset.

### 3.2 Disparity Channels

We start by modeling the retina under photopic conditions, i.e., under light conditions in which rods are saturated and cones play the main role. Retinal output is generated by three types of retinal ganglion cells: parasol, midget, and bistratified (Nassi & Callaway, 2009). Simplified, the parasol cells mainly represent intensity (luminance) while the other two represent chromatic information: the midget cells red-green colors, and the bistratified cells yellow-blue colors. We model the intensity channel, *I*, as:

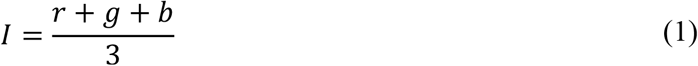

where *r, g*, and *b* are the red, green, and blue components image (Itti et al., 1998).

The color channels are modeled as below:

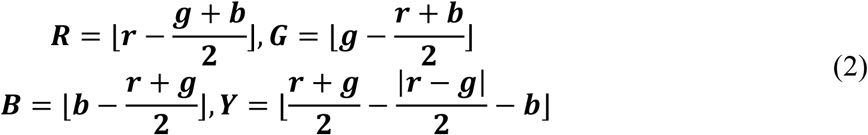

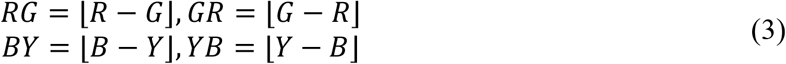

where ⌊·⌋ is half-wave rectification, and *RG, GR, BY*, and *YB* are color opponency channels. The color signals are only computed for pixels whose intensity value exceeds 10% of the maximum intensity of the input image since hue variations are not perceivable at very low luminance. It is still unclear what role chromatic information plays for stereopsis. It has been reported (Gregory, 1977; Jordan et al., 1990; Lu & Fender, 1972) that random-dot stereograms need luminance cues to cause depth perception although isoluminant figural stimuli can be perceived as stereoscopic. In our models, we implement versions with and without contributions from color channels, see below.

We use the disparity energy model as a biologically-plausible method to extract depth (Ohzawa et al., 1990, 1997). The brain, as well as our model, combines multi-spatial-frequency filters to accurately detect disparity information (Baba et al., 2015; Fleet et al., 1996; Kumano et al., 2008). The calculation of disparity energy starts by computing receptive field properties of monocular simple cells. Simple cells in V1 are modeled by Gabor filters (Kulikowski et al., 1982), in our model,

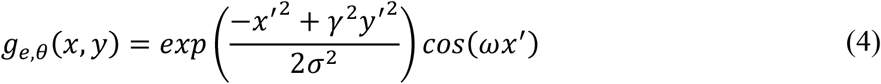

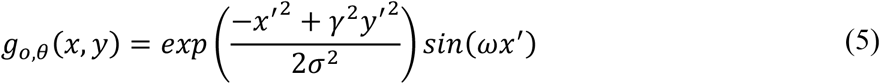

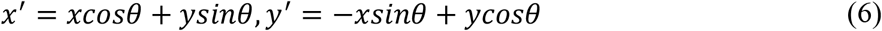

Where 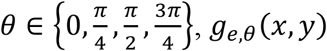 and *g*_*e, λ*_(*x, y*) are the even- and odd-Gabor filters with spatial aspect ratio *γ*, width *σ*, and spatial frequency *ω*. Frequently, the aspect ratio *γ* is chosen to be below unity, resulting in elongated filters as shown in Fig. **2**(a). For instance, Russell et al. used *γ* = 0.5 and *γ* = 0.8 for edge detection and orientation channels, respectively, and we use the same values for those purposes (edge detection and orientation channels). However, for the simple cells of the disparity features we employ shortened Gabor filters with *γ* = 5 because such filters showed better results than elongated filters. Although most orientation-selective cells in early cortex have elongated receptive fields (*γ* < 1), a fraction of them shows aspect ratios considerably greater than unity, *γ* > 1 (Xu et al., 2016). This is rare for simple cells but more common for complex cells (ibid.). An example of a shortened filter is shown in Fig. **2** (b).

Model receptive fields vary in size to make responses tolerant to changes in scale. For the sake of computational efficiency, we scale the input image, by full-octave steps, rather than the filters. The image of the *k*-th scale level is written as *β*^*k*^, *β* ∈ {*I, RG, GR, BY, YB*}. Monocular simple cells are represented as:

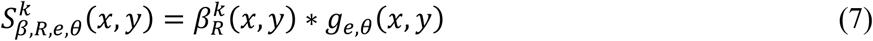

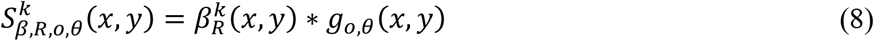

Where 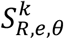 is a *k*-th level simple cell activation function with even-symmetric Gabor filters from the right image which has a preferred angle of *θ*. 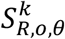 is the same but for an odd-symmetric Gabor filter. The asterisk symbol * indicates convolution. The simple cells from the left image, 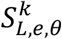 and 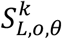, are obtained in the same way:

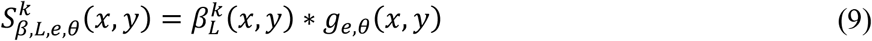

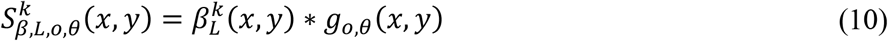

We define four 3D models which differ in the combinations of features and orientations. To compute disparity information, Model A uses intensity at one orientation (vertical), model B uses intensity at four orientations, model C uses intensity plus color features at one orientation and model D uses intensity plus color features at four orientations. This means that model A, the simplest, uses 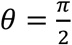 equations (4) and (5), i.e., a vertical Gabor filter and only the *I* channel as *β* in equations (7) – (10). Model D, the most complex, utilizes 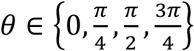 for the Gabor filters and *I, RG, GR, BY*, and *YB* channel as *β*.

Responses of binocular complex cells are calculated from the simple cells with displaced images,

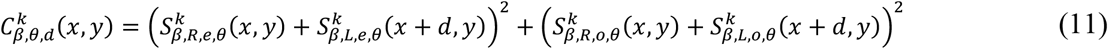

where *d* is the disparity between the right and left images. The range of the disparity *d* is arbitrary. In this paper, *d* takes on the range of ±8% of the input image width. The simple cell activities *S*^*k*^ are rescaled to the original image size before the calculation to make the disparity *d* cover the same displacement at all levels.

As described, this is called a position-based model since the displacement between two images is represented as a position difference. indicated by *d* in the equations. The displacement can also be represented by a phase difference, resulting in phase-based models (Fleet et al., 1991, 1996; Ohzawa et al., 1997). Physiological experiments show that the brain uses both approaches (DeAngelis et al., 1991; Ohzawa et al., 1990), but their roles in stereopsis are controversial. One possibility is that the phase disparity tuned cells are used because they provide higher accuracy (Qian & Zhu, 1997). However, pure phase-based disparity can only capture disparities smaller than the receptive field’s wavelength. Furthermore, Read and Cumming pointed out that phase disparity does not exist in natural images and that cells responding with phase disparity characteristics may function as “lie-detectors” to eliminate false matches (Read & Cumming, 2007). In this study, for the sake of simplicity we use only the position-based model.

The output of a binocular complex cell is enhanced when the two images match at the cell’s preferred disparity. Focusing on a specific location (*x, y*), the value of 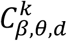 represents the “confidence” that the location belongs to the specific disparity *d*, which depends on *β, θ*, and *k*. However, its response is also enhanced where false matches or high monocular contrast exist. To compensate for such unreliable responses, we employ a softmax function to compute normalized complex cell responses *C*′,

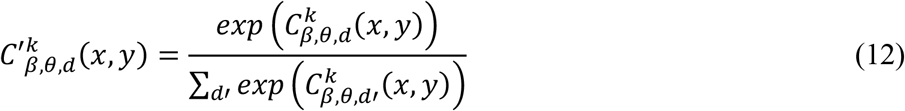

This can be interpreted as the “normalized confidence” of the disparities of each location (*x, y*) and for each of the parameters (scale, angle, color, and intensity). This computation is similar to the divisive normalization mechanism that is found in many cortical circuits (Heeger, 1992).

Then, *C*′ is linearly summed up over scales, intensity and color maps, and orientations to compute disparity confidence maps, *D*′. This is written as:

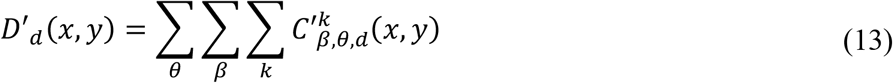

Integration of multiple spatially frequency maps was reported in visual cortex (Baba et al., 2015; Kumano et al., 2008) (although we use a broader range of frequencies in our model) as was integration of color and orientation (Garg et al., 2019; Ghose & Ts’o, 2017)

In the primate brain, disparity information is sent to both dorsal and ventral areas (Preston et al., 2008). Here, we focus on the ventral stream which encodes depth information categorically while the dorsal stream represents it in a metric manner (Preston et al., 2008). Thus, we collapse the normalized disparity map into three categories: near-, zero-, and far-positions. This can be written as:

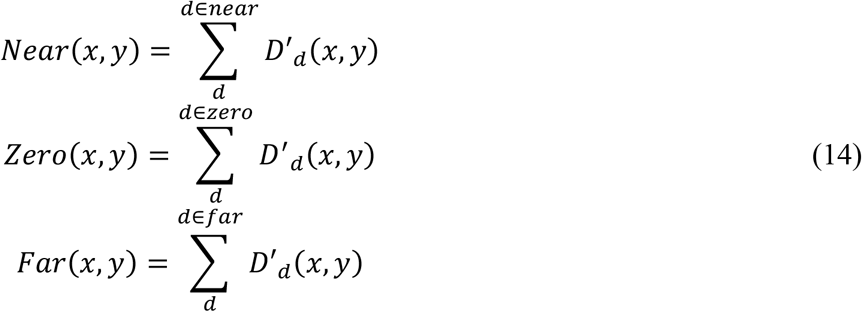

where *near, zero*, and *far* mean ranges of disparities for each position in the scene. We set these ranges based on Panum’s fusional area which is defined as encompassing any point where binocular fusion can be achieved (i.e. absence of diplopia) and which spans approximately 10 to 20 minutes of arc disparity (Qin et al., 2004). The channel of *zero* is set to approximate Panum’s fusional area, and *near* and *far* include all nearer and farther disparities, respectively. We used ±5 pixels (which corresponds to approximately ±10 min of arc for our validation setup described in Section 3.3) as the range for the *zero* channel.

Fig. **3** shows a schematic of the disparity channel computation. The calculated disparity channels form one set of inputs to the proto-object based saliency model, on the same footing with the 2D features intensity, color, and orientation (Russell et al., 2014; Uejima et al., 2020).

### 3.3 Validation

To evaluate the quality of the proposed model, we used publicly-available eye fixation datasets to compare our saliency maps with human eye movements which are taken as ground truth for the deployment of selective attention. We used the Gaze-3D dataset (Wang et al., 2013) which includes 18 images and the NCTU-3D dataset (Ma & Hang, 2015) comprising 475 images. We reduced the image size of the datasets by a factor of two before using them as input to our model to decrease computation time.

For quantitative validation, we employed five metrics to assess the predictive performance of the generated saliency maps for human fixations. The metrics are normalized scanpath saliency (NSS), Pearson’s correlation coefficient (CC), similarity (SIM), Kullback-Leibler divergence (KLD), and shuffled area under the ROC curve (sAUC). These metrics were calculated using published codes (Bylinskii et al., 2019).

As a short overview, NSS is the mean value of the normalized saliency map at the fixation locations. The normalized saliency map is calculated by transforming the map values to zero mean and unit standard deviation. CC takes on zero value for two uncorrelated variables and unit value for identical ones. The SIM measure of two maps is zero when the maps have no overlap and unity if the two maps are identical. KLD quantifies the dissimilarity between two probability distributions, and smaller KLD indicates higher similarity. The sAUC is a modified version of the area under the ROC curve. The Receiver Operating Characteristic (ROC) measures the ratio of true positives and false positives at various thresholds. The sAUC samples negative points to calculate the false positive rate from fixation locations of *other* images, rather than uniformly random locations from the *same* image that standard AUC uses. This compensates for systematic biases present in all images, such as the well-known center-bias, see. e.g. Parkhurst et al 2002 (Parkhurst et al., 2002).

It is known that blurring saliency maps can affect metrics (Borji & Itti, 2012; Hou et al., 2012). Basically, the blurring approximates the sampling error of the eye tracker used for recording fixations. We applied 2D Gaussian kernels with various widths and determined the optimal blurring kernel for each model and metric. The kernel width was varied between 1% and 20% of the image widths for NCTU-3D, and between 1% and 40% for Gaze-3D in steps of 1%. An exception was the sAUC metric which was calculated under the condition of the blurring kernel width between 1% to 8%, because the sAUC’s metric produced higher values for smaller kernels than the other metrics.

All metrics except sAUC are affected by the center-bias: human observers tend to fixate preferentially at locations in the vicinity of the centers of images. Parkhurst et al (Parkhurst et al., 2002) showed that weighting saliency with a Gaussian at the image center resulted in better fixation prediction, and that it could be improved even more by centering a Gaussian on the location of the instantaneous fixation, to take into account the fall-off of visual acuity in the periphery. Following (Zhang & Sclaroff, 2016), here we use a simpler approach of using a fixed parabolic distance-to-center (DTC) re-weighting,. This is computed as:

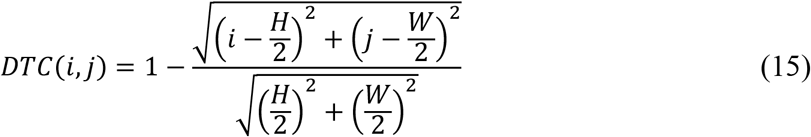

where *i* and *j* are the row and column index and *H* and *W* are the height and width of the input image. The generated saliency map blurred by the 2D Gaussian kernel with the best sigma is pixel-wise multiplied with the DTC. The DTC re-weighting procedure was not applied for the sAUC metric because sAUC automatically compensates for the center-bias.

## 4 Results

Examples of the proposed disparity channels and the resulting saliency map are shown in Fig. 1 (b) and (c), respectively. The fixation map in this example indicates that participants tend to fixate a fish in the lower half of the image that is closer to the observer than the other fish. The computed disparity maps in Fig. 1 (b) show that the location of the fish is captured by the near and zero channels. The saliency map is calculated based on the disparity channels and shows high value at the approximate location of this fish as shown in Fig. 1 (c)

Fig. 4 shows comparisons between fixation maps and saliency maps generated by the 2D and 3D models. We here use model A described in Section 3.2 which employs vertical orientation and intensity. The 2D model saliency maps were calculated from intensity, color, and orientation features as in (Uejima et al., 2020), but without texture features. The 3D saliency maps were computed by adding depth maps generated from the disparity channels. In these examples, the 2D model predicted the human fixations to some extent, and the depth information improved the predictions. In the example of the first row, for instance, the 3D model shows higher saliency value at the location of the airship than the 2D model, in agreement with human fixations. In all examples shown, the 3D model suppresses saliency to background patterns, relative to foreground objects. This shows that depth plays a role in the prediction of overt attention.

**Fig. 4.**
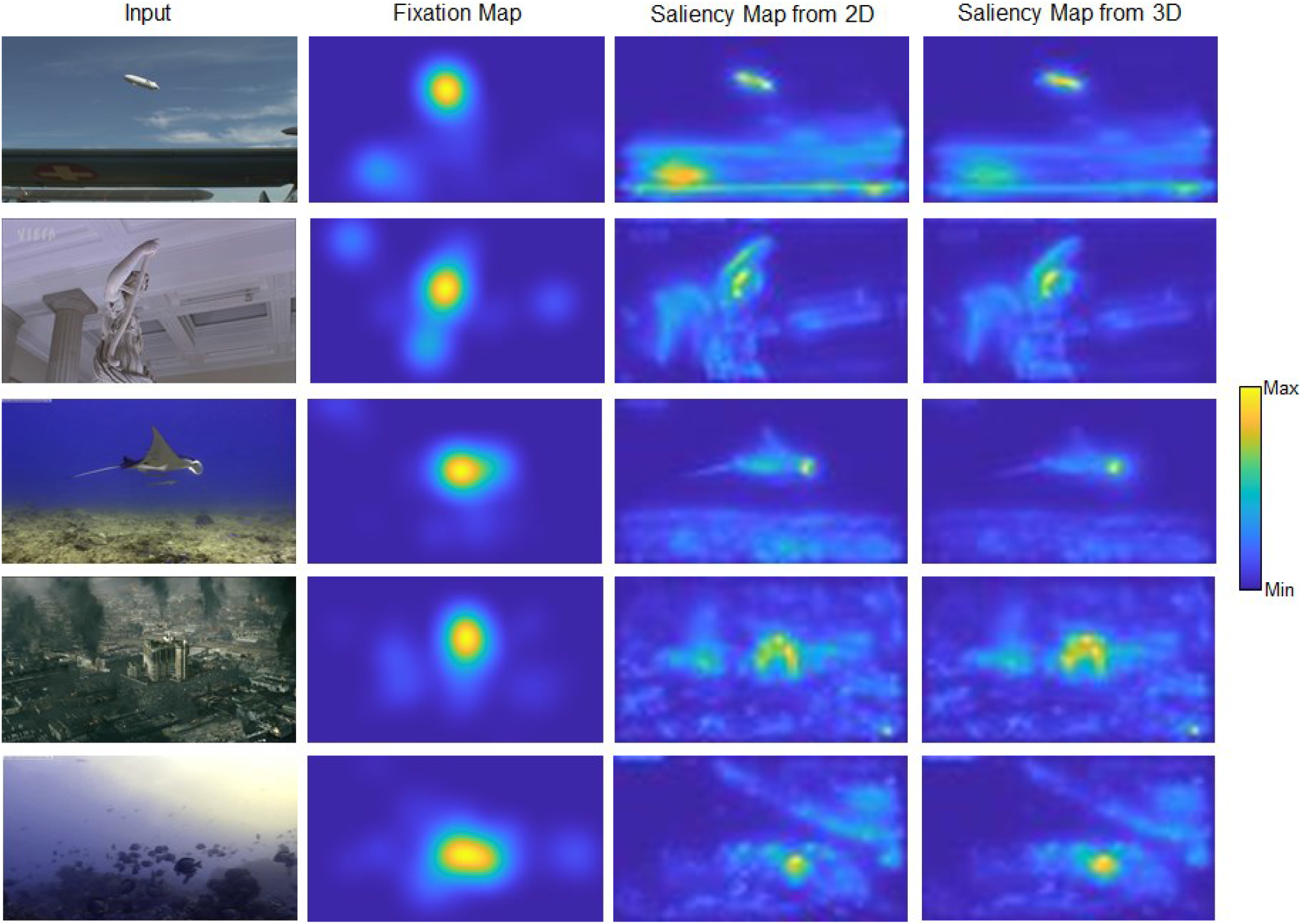
Examples of saliency maps generated by the 2D model and a 3D model (model A). The input image and fixation map are from the NCTU-3D dataset. Our 3D model suppresses background saliency compared to the 2D model.

To quantify the performance enhancement due to adding depth channels, we calculate the metrics of the saliency maps generated with and without the depth information by comparing them against human fixations. These metrics are shown in Table 1. As described in Section 3.2, we used four variations of the 3D model which include different combinations of color features and orientations. Our results indicate that the depth channels improve the prediction of human fixations. Because of the small size of the Gaze-3D dataset (only 18 images), in the following we focus our analysis on the NCTU-3D data.

**Table 1.**
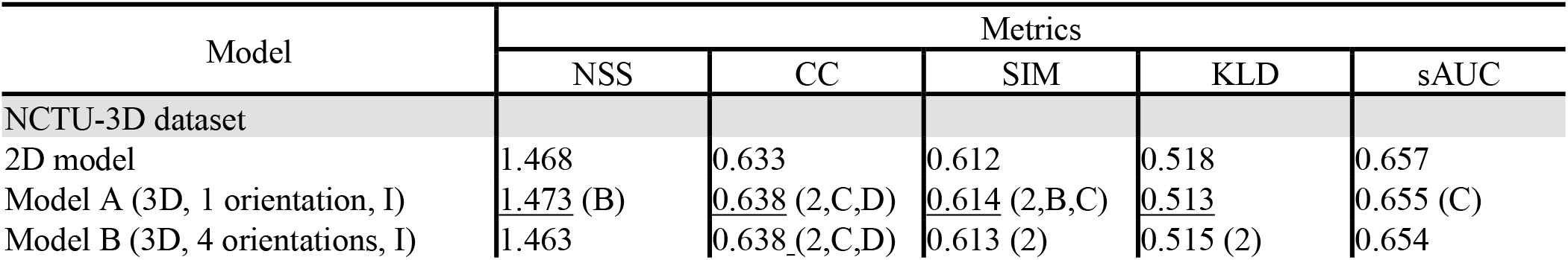
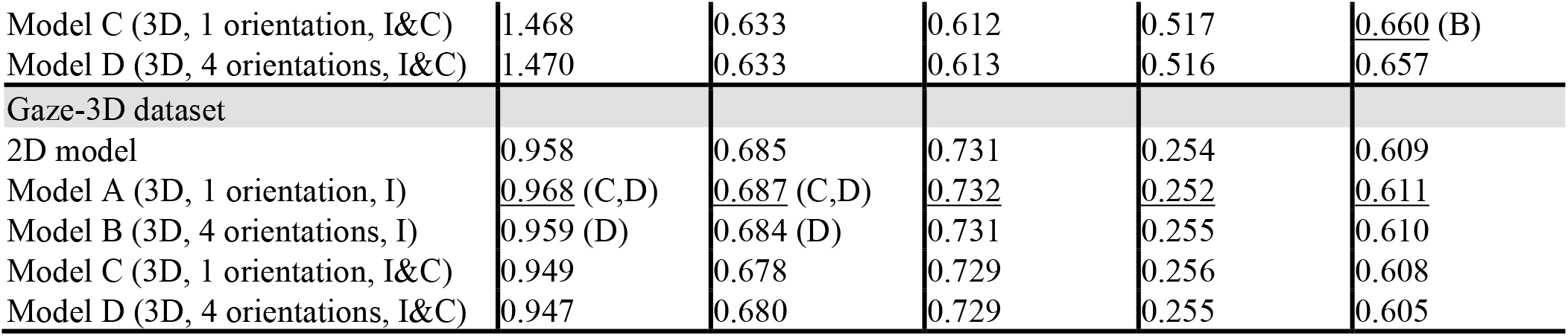
Models incorporating depth features predict human eye fixations equally well or better than the 2D model. In column 1, “1 orientation” indicates a model with only one (vertical) Gabor filter and “4 orientations” models with Gabor filter with four orientations. Labels “I” and “I&C” indicate models with only the intensity feature and a combination of intensity and color features, respectively. Underscore denotes the best score for each metric. A parenthesis next to the performance value of a model indicates that this model performs significantly better than any of the models listed in the parenthesis, where B, C, D, indicate the different 3D models, and “2” the 2D model. For instance, by the CC metric Model A performs better than models 2, B, and C. Significance was evaluated by two-tailed paired t-tests (*p* < 0.05). Larger values are better for all metrics but KLD.

| Model | Metrics |  |  |  |  |
| --- | --- | --- | --- | --- | --- |
|  | NSS | CC | SIM | KLD | sAUC |
| NCTU-3D dataset |  |  |  |  |  |
| 2D model | 1.468 | 0.633 | 0.612 | 0.518 | 0.657 |
| Model A (3D, 1 orientation, I) | <u>1.473</u> (B) | <u>0.638</u> (2,C,D) | <u>0.614</u> (2,B,C) | <u>0.513</u> | 0.655 (C) |
| Model B (3D, 4 orientations, I) | 1.463 | <u>0.638</u> (2,C,D) | 0.613 (2) | 0.515 (2) | 0.654 |
| Model C (3D, 1 orientation, I&C) | 1.468 | 0.633 | 0.612 | 0.517 | <u>0.660</u> (B) |
| Model D (3D, 4 orientations, I&C) | 1.470 | 0.633 | 0.613 | 0.516 | <u>0.657</u> |
| Gaze-3D dataset |  |  |  |  |  |
| 2D model | 0.958 | 0.685 | 0.731 | 0.254 | 0.609 |
| Model A (3D, 1 orientation, I) | <u>0.968</u> (C,D) | <u>0.687</u> (C,D) | <u>0.732</u> | <u>0.252</u> | <u>0.611</u> |
| Model B (3D, 4 orientations, I) | 0.959 (D) | 0.684 (D) | 0.731 | 0.255 | 0.610 |
| Model C (3D, 1 orientation, I&C) | 0.949 | 0.678 | 0.729 | 0.256 | 0.608 |
| Model D (3D, 4 orientations, I&C) | 0.947 | 0.680 | 0.729 | 0.255 | 0.605 |

We first look at the models that only use intensity information (not color), i.e., models A and B. For all but two of the ten comparisons between the 2D model and the two corresponding models with 3D information (Models A and B), the models with 3D information performed equally well or better than the 2D model. The increase in performance was significant for 5 of these comparisons (two-tailed paired t-test, p < 0.05).

If color information is added, models incorporating 3D cues (Models C and D) are equal or better than the corresponding 2D model for all ten comparisons. However, the differences are small and none of them reaches significance.

Primates express strong interest in faces and bodies which attract attention even when they are task-irrelevant (Landman et al., 2014). Indeed, detection of faces and body parts is supported by anatomical structures in monkeys (Desimone, 1991; Gross, 2008; Tsao et al., 2003) and humans (Downing et al., 2001). We expect that the human fixation locations that we use as ground truth in this study show a similar bias. Since none of our models has corresponding explicit detection mechanisms for faces or body parts, we expected that models predict fixation locations better for images that have no persons in the scene than for images with persons. We therefore divided the images into two sets: one set with humans visible (fully or partially) and the other set without. The latter could, however, include animals or statues of humans. In the NCTU data set, we found 303 images in which humans were visible, at least partially, and 172 images in which that was not the case. In the Gaze-3D data, 7 images included visible humans and 11 did not.

The results shown in Table 2 confirm our expectation: for all five models, and for all five metrics, fixation prediction performance decreased with the presence of faces, bodies, or body parts in the NCTU-3D dataset. In the majority of the tests (15/25), this decrease was significant (one-tailed Welch test, p<0.05). Similarly for the Gaze-3D data, the majority (17/25) of tests showed higher performance for images without humans and many of these differences (10/17) was significant. Overall, the effect was somewhat weaker than for the NCTU-3D data, but the Gaze-3D dataset is very small which limits the statistical power that can be achieved. We also note that a separate channel for face detection, for instance using the standard Viola-Jones algorithm (Viola & Jones, 2001), can be easily added to the models and would most likely increase fixation prediction performance substantially, as it did in a previously-developed class of models from the same pedigree as ours (Cerf et al., 2008).

**Table 2.** Effect of presence of persons in images. The left and right columns of each metrics show model performance on images without and with persons, respectively. Underscore indicates that the performance is better on images without persons. For the NCTU-3D data, model performance is always higher on images without persons. This is not the case for Gaze-3D but this dataset has only 7 images with persons. An asterisk (*) indicates statistical significance (one-tailed Welch test, *p* < 0.05).

| Model | Metrics |  |  |  |  |  |  |  |  |  |
| --- | --- | --- | --- | --- | --- | --- | --- | --- | --- | --- |
|  | NSS |  | CC |  | SIM |  | KLD |  | sAUC |  |
| NCTU-3D dataset | without | with | without | with | without | with | without | with | without | with |
| 2D model | <u>1.540</u> | 1.427 | <u>0.685</u>* | 0.603 | <u>0.637</u>* | 0.597 | <u>0.478</u>* | 0.541 | <u>0.657</u> | 0.657 |
| Model A (3D, 1 orientation, I) | <u>1.551</u>* | 1.430 | <u>0.693</u>* | 0.607 | <u>0.640</u>* | 0.599 | <u>0.465</u>* | 0.541 | <u>0.661</u> | 0.653 |
| Model B (3D, 4 orientations, I) | <u>1.545</u>* | 1.416 | <u>0.692</u>* | 0.607 | <u>0.640</u>* | 0.598 | <u>0.469</u>* | 0.542 | <u>0.655</u> | 0.653 |
| Model C (3D, 1 orientation, I&C) | <u>1.532</u> | 1.432 | <u>0.685</u>* | 0.604 | <u>0.635</u>* | 0.599 | <u>0.479</u> | 0.539 | <u>0.665</u> | 0.657 |
| Model D (3D, 4 orientations, I&C) | <u>1.543</u> | 1.429 | <u>0.685</u>* | 0.604 | <u>0.637</u>* | 0.599 | <u>0.476</u> | 0.538 | <u>0.660</u> | 0.654 |
| Gaze-3D dataset | without | with | without | with | without | with | without | with | without | with |
| 2D model | 0.874 | 1.090 | <u>0.703</u> | 0.657 | <u>0.760</u>* | 0.686 | <u>0.217</u>* | 0.311 | <u>0.610</u> | 0.607 |
| Model A (3D, 1 orientation, I) | 0.878 | 1.110 | <u>0.704</u> | 0.660 | <u>0.760</u>* | 0.690 | <u>0.219</u>* | 0.304 | <u>0.618</u> | 0.599 |
| Model B (3D, 4 orientations, I) | 0.871 | 1.098 | <u>0.702</u> | 0.656 | <u>0.759</u>* | 0.687 | <u>0.222</u>* | 0.307 | 0.604 | 0.621 |
| Model C (3D, 1 orientation, I&C) | 0.855 | 1.096 | <u>0.693</u> | 0.655 | <u>0.757</u>* | 0.684 | <u>0.220</u>* | 0.312 | 0.606 | 0.610 |
| Model D (3D, 4 orientations, I&C) | 0.862 | 1.082 | <u>0.696</u> | 0.653 | <u>0.758</u>* | 0.684 | <u>0.216</u>* | 0.316 | 0.598 | 0.616 |

## 5 Discussion

Binocular information processing in our model is based on physiological and psychological evidence. Its basic mechanism is the binocular energy model which combines information from two monocular input sources into a binocular signal, akin to the generation of complex cell responses from the activity of monocular simple cells in two eyes. Binocular complex cells are tuned to specific disparity ranges and their activity represents a “confidence” measure of the disparity difference within their receptive fields. Our model of cells with binocular receptive fields uses Gabor filters with low aspect ratio (<1). While the aspect ratio of the majority of orientation selective cortical cells is >1, a fraction of cells in early visual cortex have an aspect ratio smaller than unity. We hypothesize that their role is primarily in disparity computations, rather than in the representation of oriented edges for which elongated filters are better suited.

Despite decades of neurophysiological research, it is still not entirely clear how the brain deals with depth information. Early studies of stereopsis proposed that the disparity processing is achieved categorically, by cells tuned near, far, and zero (in the focal plane) (Poggio & Poggio, 1984; Richards, 1971). Later, this three-channel model was replaced by a continuous representation, similar to that for orientation or motion direction (Poggio, 1995). A recent imaging study indicates that, in fact, both ideas may be correct but used in different pathways: dorsal cortical areas, including V3A and V7encode metric disparity while areas in the ventral pathway, in particular the lateral occipital area, represent the categorical responses “near” and “far (Preston et al., 2008). The former may be more relevant for visual control of action while the latter may be more useful for tasks like object recognition. In our model, we adopt the latter, i.e. a categorical model akin to what is found in the ventral pathway, with the responses of the binocularly tuned cells organized in three channels: near, far, and (close-to-)zero.

The boundaries between the zero channel and the other two are defined by Panum’s area. This gives rise to a limitation of our approach because the model assumes that the focal plane is always in the zero disparity range of the input image. Because subjects can move their eyes, they can fixate objects outside the focal plane in which case Panum’s fusion area changes. Our model does not include such dynamic change of subjective disparity perception.

The output of these disparity channels is given as input to a proto-object based saliency model (Russell et al., 2014) which means that proto-objects are calculated separately in the spaces of near, far, and zero channels. As in earlier saliency map models, interactions between features and spatial scales emphasize the influence of those maps with a small number of local maxima and suppress those with many peaks. In typical scenes, a small number of foreground objects tend to be in the near or zero depths while broad areas of cluttered background are in the “far” channel. For such scene structures, the model emphasizes the foreground objects in the near and zero channels which only have a few peaks. This agrees with the observation that humans tend to fixate objects in foreground (near) locations (Gautier & Le Meur, 2012; Jansen et al., 2009; Lang et al., 2012). We find that by most metrics our intensity-based models (Models A and B) predict human fixation patterns better than the analogous model that uses monocular information only (Table 1).

This was not the case for the two models that, in addition to intensity, also use color information. While intensity-only based Models A and B showed nearly uniformly better performance than the 2D model, the models incorporating color information to calculate depth in addition (Models C and D) performed similarly to the 2D model on the NCTU-3D dataset, and even slightly worse on the Gaze-3D dataset (the small size of Gaze-3D makes it difficult to assign high importance to the latter result). Furthermore, comparing the intensity-only models directly with the models that use both intensity and color information, the former predict fixation clearly better. This result seems at first puzzling: why would a model that has access to more information perform worse than one with less information?

There are two considerations to take into account. First, parameters in our models are assigned fixed values, they are not selected by a learning algorithm that optimizes performance in a given task (here, optimal agreement with eye fixations). There is therefore no guarantee that providing additional information improves the performance. Second, and in addition, we define performance as agreement with human eye movements. By this measure, the expectation that making available additional source of types of information improves performance rests on the assumption that this information is also available to, and used by, the mechanisms that guide eye movements in humans. This is, however, not necessarily the case; the role of chromatic information in stereopsis is complicated. At the same time that Julesz invented random-dot stereograms (Julesz, 1971), he reported that anti-correlated dots (with reversed intensity polarity) did not give rise to stereoscopic vision, but that correlated colors of the dots aided binocular fusion. Shortly thereafter, Lu and Fender disputed the latter finding, concluding their paper with the sentence “luminance alone is used as the principal signal to determine their depth” (Lu & Fender, 1972). Later studies found evidence for stereoscopic vision in isoluminant conditions (Comerford, 1974; Grinberg & Williams, 1985) but methodological problems with their approach led Livingstone and Hubel to question these results (Livingstone & Hubel, 1987). Instead, they proposed that color, processed in the parvocellular stream, interacts minimally, at best, with orientation, motion and depth which are predominantly represented in the magnocellular stream. This interpretation has been questioned yet again by newer results that provide evidence for effective cross-communication between the magnocellular and parvocellular channels that support stereopsis driven by luminance, even though it is not clear that color information by itself is sufficient to induce depth perception (Scharff & Geisler, 1992; Simmons & Kingdom, 1997; Tyler & Cavanagh, 1991). While newer evidence supports that color information does modulate perception of depth (Den Ouden et al., 2005), it is not clear if the effect is strong enough in natural scenes, the stimuli that we use, to result in statistically significant differences. Overall, while it was not the focus of our work to study the role of color in stereopsis, our results provide some support to the notion that for this perceptual function and for this set of stimuli, chromatic information pays only a minor role if any.

Understanding stereoscopic vision is not only of interest for basic science but it also has practical implications. As mentioned in the Introduction, one is obviously the determination of the distance of objects from the observer. Another (related) one is camouflage breaking. The goal of camouflage is to disrupt the process of segmenting the to-be-camouflaged object from its background. In addition to the obvious method of trying to match the local visual properties of the object as closely as possible with those of the background, a (related) technique is to create strong contours inside the object boundaries, resulting in the creation of internal “false” edges. This process interferes with the grouping of the object’s features into a coherent entity, and therefore its identification (Adams et al., 2019). Availability of depth information reduces the effect of monocular edges internal to the object and therefore aids the visual system in the formation of correct object borders, segregating the object from the background. It is known that visual scenes are organized in terms of such segregated objects, or more precisely, proto-objects (Rensink, 2000) which are entities towards attention is directed (Egeth & Yantis, 1997; Egly et al., 1994; Scholl, 2001). The fact that our 3D models predict human fixation patterns better than the 2D model supports the idea that human fixations are partially guided by depth cues. Furthermore, this result may have consequences for the application of our models in the context of camouflage. Given that depth cues contribute strongly to scene segmentation and, therefore, camouflage breaking (Adams et al., 2019), we may conjecture that this is a possible application where algorithms like our models A and B may be useful for this purpose. We did not test this hypothesis specifically on scenes with camouflage patterns, but this is a topic for future research.

## 6 Conclusion

We incorporate a biologically plausible stereopsis mechanism into a proto-object based saliency model. The proposed model takes stereoscopic images as input and computes categorical depth information from a disparity energy model. We combine the resulting depth information with monocular information (intensity, color, and orientation) to form a representation of the visual scene in terms of proto-objects. The resulting saliency map generates predictions for the allocation of overt attention that agree significantly better with human behavior than those from the corresponding maps using monocular cues only. We also note that, different from typical machine learning approaches, all stages of this process are derived from first principles. No training is required other than the setting of a small number of general parameters

## Acknowledgments

We thank the National Science Foundation and the National Institutes of Health which support this work through the CRCNS and the NCS-FO mechanisms. T. Uejima’s PhD training was supported by the Japanese Acquisition, Technology & Logistic Agency, Government of Japan.

